# E.PAGE: A curated database and enrichment tool to predict gene modules associated with gene-environment interactions

**DOI:** 10.1101/2022.01.03.474848

**Authors:** Sachin Muralidharan, Sarah Ali, Lilin Yang, Joshua Badshah, Farah Zahir, Rubbiya Ali, Janin Chandra, Ian Frazer, Ranjeny Thomas, Ahmed M. Mehdi

**Author notes:** Address correspondence to Dr. Ahmed Mehdi, The University of Queensland Diamantina Institute, Translational Research Institute, 37 Kent St, Woolloongabba, Qld, 4102, Australia. equal contribution (Sachin Muralidharan, Sarah Ali, Lilin Yang). Senior authors (Ian Frazer, Ranjeny Thomas, Ahmed M. Mehdi).

## Abstract

**Background:** The purpose of this study was to manually and semi-automatically curate a database and develop an R package that will provide a comprehensive resource to uncover associations between biological processes and environmental factors in health and disease.

We followed a two-step process to achieve the objectives of this study. First, we conducted a systematic review of existing gene expression datasets to identify those with integrated genomic and environmental factors. This enabled us to curate a comprehensive genomic-environmental database for four key environmental factors (*smoking, diet, infections* and *toxic chemicals*) associated with various autoimmune and chronic conditions. Second, we developed a statistical analysis package that allows users to interrogate the relationships between differentially expressed genes and environmental factors under different disease conditions.

**Results:** The initial database search run on the Gene Expression Omnibus (GEO) and the Molecular Signature Database (MSigDB) retrieved a total of 90,018 articles. After title and abstract screening against pre-set criteria, a total of 186 studies were selected. From those, 243 individual sets of genes, or “gene modules”, were obtained. We then curated a database containing four environmental factors, namely *cigarette smoking, diet, infections* and *toxic chemicals*, along with a total of 25789 genes that had an association with one or more of these gene modules. In six case studies, the database and statistical analysis package were then tested with lists of differentially expressed genes obtained from the published literature related to type 1 diabetes, rheumatoid arthritis, small cell lung cancer, COVID-19, cobalt exposure and smoking. On testing, we uncovered statistically enriched biological processes, which revealed pathways associated with environmental factors and the genes.

**Conclusions:** A novel curated database and software tool is provided as an R Package. Users can enter a list of genes to discover associated environmental factors under various disease conditions.

## Background

Organisms are constantly being exposed to a wide range of environmental triggers that influence gene expression, resulting in several diseases. Environmental factors, such as drugs, toxic chemicals, smoke, temperature, dietary components and infections are considered modifiable causes of disease through their effects on biological processes, and in response, the expression of many genes is altered (1). It is estimated that environmental factors account for approximately 70% percent of all autoimmune diseases and 80% of all chronic diseases (2). These large proportions indicate that environmental exposures are an important contributor to disease, and there is ample evidence to support complex interrelationships between various environmental and genomic factors for disease causation (3). Manipulation of environmental triggers and the host immune system during the clinical and preclinical stages of a disease will offer significant insight and guide early intervention for many disorders (4).

In the era of Big Data technologies, several genomic databases exist to explore differential expression of genes under various clinical conditions(5, 6). However, to our knowledge there is currently no computational tool that can use information from existing large-scale databases to predict gene-environment relations. Therefore, in this study we formulated an integrated and comprehensive database that will provide insights of how environmental factors are associated to gene expression and disease, and leading to the identification of potential therapeutic strategies for the prevention and control of diseases attributable to both environmental and genetic factors.

## Implementation

We followed a two-step approach to conduct this study. First, we conducted a systematic review using a standard approach to identify all studies that used integrated datasets containing comprehensive information about environmental and genetic risk factors for various diseases. Second, we curated a database and developed a statistical analysis package to enable the user to understand the relationships between differentially expressed genes and select environmental factors.

### Step 1: Systematic review

The aim of this step was to identify the relevant published literature from where we could obtain existing data pertinent to gene expression changes in response to an environmental factor. In detail the systematic review was conducted as follows:

#### Search strategy

We undertook a comprehensive literature and database search using PubMed, Gene expression omnibus (GEO), and Gene set enrichment analysis (GSEA) databases (7). All databases were searched from their inception until 16th October 2020. The reference lists of all the retrieved studies were examined to identify additional studies.

The search terms and their synonyms related to environmental factors and gene expression. The keywords used included medical subject headings (MeSH) terms, e.g., (“Diet”[MeSH Terms] OR diet [All Fields]) AND (“gene expression”[MeSH Terms] OR gene expression [All Fields]). Table 1 details the search strategy and date of searches for various databases.

**Table 1.** Search strategies used for database searching.

| Search term | Number of hits (Total) | Date of search hits |
| --- | --- | --- |
| Cigarette smoking AND Gene expression | 324 | 16/10/2020 |
| Diet AND Gene expression | 25440 | 16/10/2020 |
| Infection AND Gene expression [GEO Database] | 59338 | 16/10/2020 |
| C7 Immunologic gene sets [GSEA] | 4872 | 16/10/2020 |
| Toxic chemical AND Gene expression | 44 | 16/10/2020 |

Inclusion/exclusion criteria:

Pre-set inclusion criterion for studies to be considered eligible were:

- Only articles written in English
- Participants of any age group and both genders.
- Since most of the experimental trials involving environmental factors were carried out in humans or mice, we included hits for Homo sapiens and Mus musculus.
- Four specific environmental factors were chosen, based on the previous published evidence for major contribution as an environmental factor affecting gene expression (8). Specifically,
  - Cigarette smoking – Includes data related to the practice of tobacco smoking and inhalation of tobacco smoke.
  - Diet - Includes data on the various types and quantities of food consumed by a person.
  - Infections - Includes data on infections caused by pathogenic organisms such as viruses, bacteria, fungi, protozoa and parasites.
  - Toxic chemicals - Includes data on substances such as metals or other chemical agents that are hazardous to human health if inhaled, ingested or absorbed.
- We included published data from datasets, series and platforms. Samples were excluded if they consisted of unpublished data. We did not limit the search specific for any disease.

We did not include any dataset relating to mRNA, protein, CDS or small non-coding RNAs like miRNA or siRNA.

#### Literature review method

Two reviewers screened the abstracts and citations independently at the same date and time and using the same search parameters. We identified articles that met the inclusion criteria. After title and abstract screening, studies were selected for full-text review. After the full length article review, those studies that met the inclusion criteria were selected for data extraction (7).

#### Data extraction

Two reviewers independently extracted data. The specific features extracted from each article were: (1) Differential gene expression data; (2) specific description of the type of data collected; (3) specific keywords related to the differentially expressed genes for each dataset, including disease, sample condition and pathways. These were manually searched in the abstract, demographics and result sections of each publication.

Data were extracted and coded in a spreadsheet to collate information from each study. The data were combined and any anomalies between reviewers were resolved by a third reviewer.

#### Quality and data validity assessment

The methodological quality was checked before including the data, using the Q-Genie tool (9). We recorded whether the study used a standard microarray procedure and descriptions of the sample data, causes of up- and downregulation of genes and any other specific changes in the gene expression.

### Step 2: Software generation

The statistical analysis package E.PAGE (Environmental Pathways Affecting Gene Expression) (github.com/AhmedMehdiLab/E.PAGE) was written in R version 4.0.3 (www.R-project.org/) and developed using RStudio (rstudio.com). Using publicly available packages (tidyverse www.tidyverse.org, Seurat satijalab.org/seurat/) as dependencies, the package performs enrichment analysis as previously described by Mehdi and colleagues (10).

For the enrichment analysis of gene modules, we followed standard methods to perform gene set enrichment analysis. We compared the number of genes that had a specific gene modules against those that did not. A hypergeometric distribution was used to determine a p-value, which was corrected using false discovery rate (FDR) for multiple hypothesis testing using the Benjamini and Hochberg correction method (Hochberg, 2018). The results are filtered based on the adjusted P value, and those with P ≤ 0.05 are displayed to the user. Fold enrichment was calculated by taking the ratio of a set of genes containing a specific gene modules, and the total set of genes was obtained by taking the union of all the collected gene modules (10). Adjusted fold enrichment was measured as a ratio of the fold enrichment value to the negative log of adjusted P value. This log fold change value was used to determine the significance of the enriched genes specific for an gene module (10). An odds ratio then was measured to determine the probability of finding the set of enriched genes specific to an gene module (Szumilas, 2010).

### Step 3: Case studies

We used six case studies to test our enrichment tool, these studies were not used in database curation. Case study 1 involves gene expression data in peripheral blood mononuclear cells (PBMC) in children with type 1 diabetes (11). Gene expression changes were identified using microarray analysis from 43 patients with new onset T1D compared with 24 healthy controls. The gene expression data set in case study 2 is taken from the GEO database (microarray datasets; GSE12021, GSE55457, GSE55584 and GSE55235) that includes samples from 45 patients with rheumatoid arthritis, compared with 29 healthy control samples (12). Case study 3 includes gene expression data from 23 small cell lung cancer samples and 42 healthy lung tissues (13). The gene expression data from the case study 4 was taken from cobalt-exposed rat liver derived cells (14). The final two case studies used differentially expressed genes extracted from single-cell expression data. Case study 5 was based on single-cell RNAseq data from COVID-19 patients, comparing severe and healthy cases in peripheral immune environments (15), while case study 6 was based on a single-cell RNAseq-based atlas of epithelial cell-specific responses to smoking (16). For single-cell RNA seq data, E.PAGE used a Seurat object (with clustering performed) as an input and performs differential expression analyses between the clusters to uncover lists of genes to compute related enriched gene modules.

## Results

### Systematic review and E.PAGE structure

The initial electronic search of GEO and MSigDB database identified a total of 90,018 studies (Figure 1). Title and abstract screening of retrieved studies resulted in a total of 3547 studies which had potential data related to environmental factors. After full text examination of 3547 studies, 3008 studies were excluded since they did not provide any differential gene expression data associated with any of the four environmental factors. A total of 243 datasets were obtained from 186 studies and the gene expression data were retrieved and collated to form a database. Figure 1 illustrates a flow chart of all the steps taken to obtain the data that satisfy the required parameters. The overall structure of E.PAGE is shown in Figure 2. After literature screening, a database of 243 datasets was developed by linking each dataset with published lists of differentially expressed genes and the gene modules. Specifically, the text of these 186 publications and associated datasets were manually screened to develop gene modules representing the type of experiment, experimental conditions or disease type, experimental factors, demographics of subjects, and published pathways as previously described by Mehdi and colleagues (10). The final database consisting of 243 datasets is obtained through GEO and MSigDB databases and includes 18015 genes for *diet*, 13259 genes for *infections*, 3841 genes for *cigarette smoking* and 644 genes for *toxic chemicals*.

**Figure 1.**
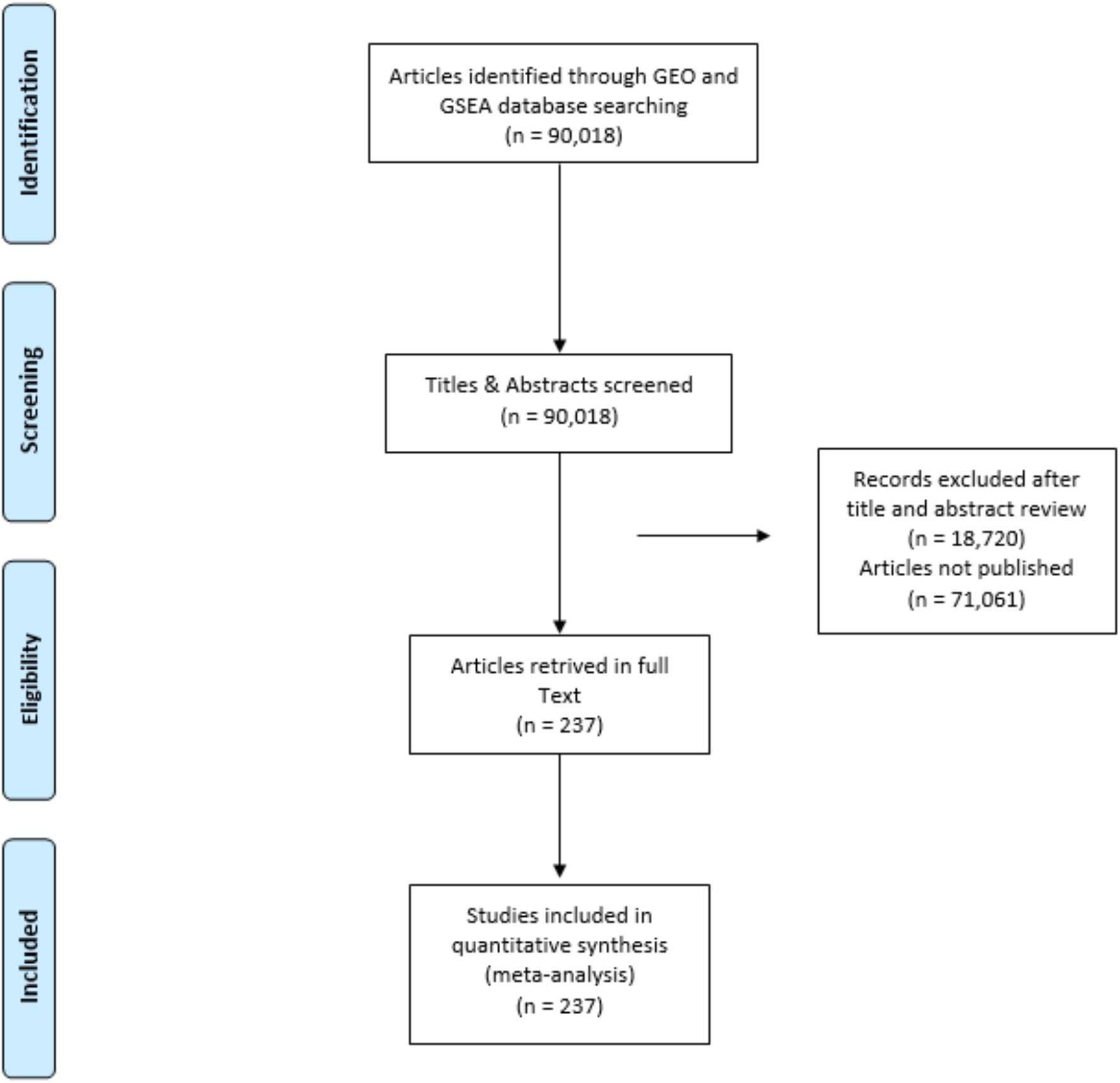
PRISMA Flow chart representing the various stages of screening involved in the systematic review process.

**Figure 2.**
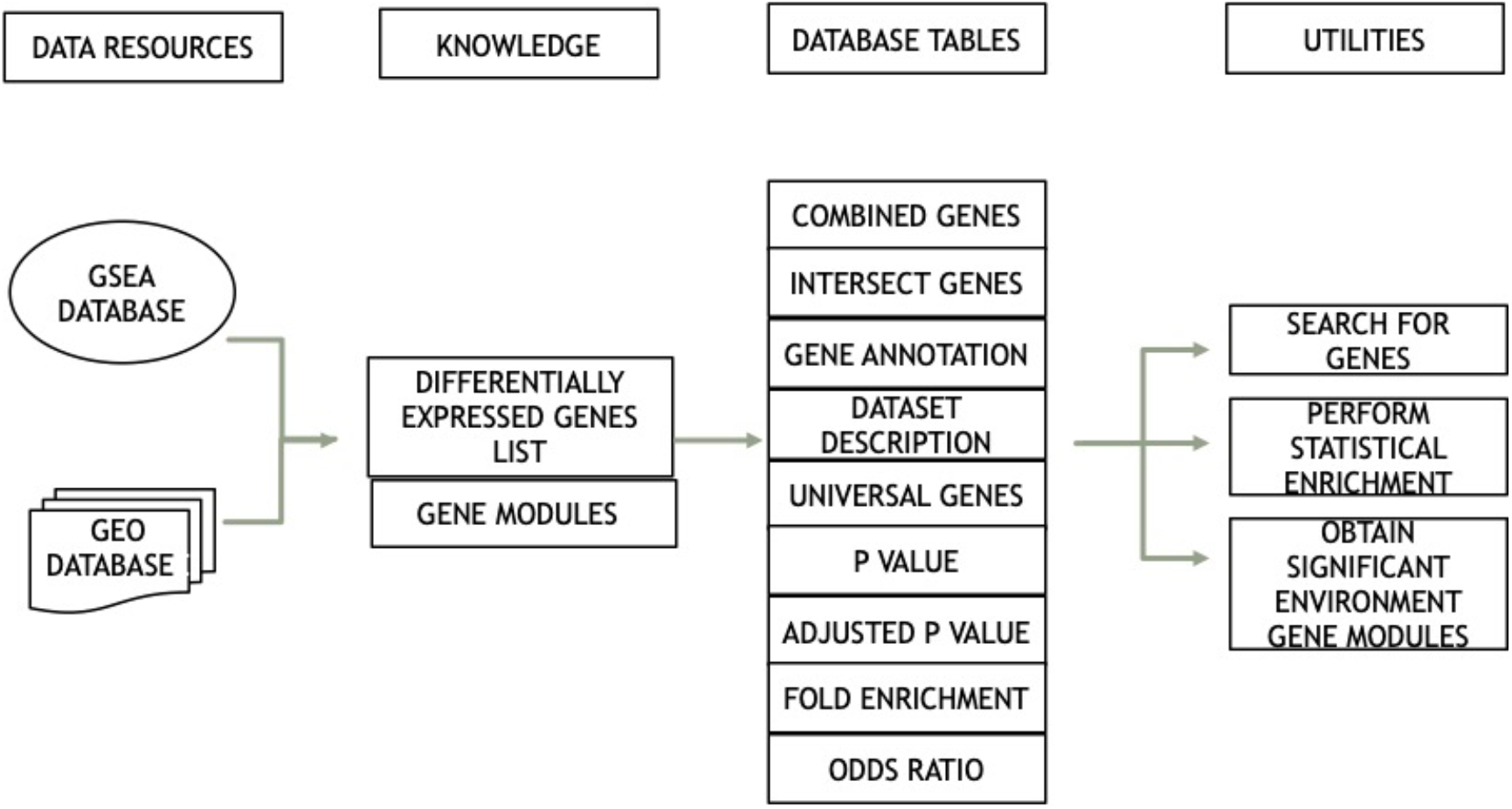
Flow chart representing the various parameters and their utilities provided on database query.

### Querying E.PAGE

An R package was developed to enable statistical enrichment and gene modules associated with datasets/genes of interest to a user. The package produces various data tables as shown in Figure 2 and a user can search genes of interest for their statistical enrichment. To test the utility of the statistical analysis package, we performed six case studies as described hereafter.

### Case studies 1 and 2: Immune response activation in type-1 diabetes and rheumatoid arthritis

We first tested whether query signatures associated with T1D and RA could recover common pathways associated with these autoimmune disease. We used 291 DE genes uncovered from 43 patients with new-onset T1D as compared to 24 healthy controls (8) (Table 2) and 229 DE genes from 45 samples from patients with RA, compared with 29 healthy control samples (12) (Table 3). The statistical enrichment using E.PAGE identified that the genes in both datasets are involved in *Immune response*. Other significant gene modules that were common to both diseases include *Interferons, IL-12* and *Transcription regulation*. These processes are all well known to be involved in RA and T1D (17). *Insulin resistance* and *Xenobiotic metabolism*, which are both believed to be associated with T1D, were uncovered using E.PAGE and validate the utility of the platform (Table 2). Similarly, for RA, many smoking related gene modules such as Smoking history and Pack years (*Smoking Status: Current, Never, Pack-years: (10 - 20), Pack-years: (20 - 30; Healthy smoker), (Above 40; Smoker with COPD)*), were uncovered indicating an important risk factor for this disease (Table 3). For both T1D and RA, a large number of gene modules related to infections, both viral and bacterial (*Lyme disease, Borrelia burgdorferi, HBV Infection, Viral response, Bacterial infection, Zika virus, Influenza A Infection, HIV infection, Echovirus-30, Rhinovirus infection*), were significantly associated with disease, indicating that similar responses are occurring in patients suffering from these chronic autoimmune diseases as in responses to infections.

**Table 2.** Collation of results obtained on query of E.PAGE with genes differentially expressed in Type 1 Diabetes.

| Gene Modules | Number of Modules | Number of DE Genes | Padj | Fold Enrichment |
| --- | --- | --- | --- | --- |
| Dyslipemia | 1 | 13 | 1.46E-08 | 12.88 |
| Olive oil induced gene expression | 3 | 15 | 7.52E-08 | 8.58 |
| Diet intake: Olive oil | 2 | 14 | 1.14E-07 | 8.96 |
| Inflammation | 31 | 82 | 6.87E-07 | 1.85 |
| Infection type: Acute | 58 | 112 | 7.46E-07 | 1.61 |
| Transcription regulation | 11 | 33 | 8.00E-07 | 3.09 |
| Interferons | 15 | 53 | 1.66E-06 | 2.22 |
| IL-12 | 4 | 37 | 5.02E-06 | 2.60 |
| Th1-mediated response | 4 | 37 | 5.02E-06 | 2.60 |
| Parasite killing | 4 | 37 | 5.02E-06 | 2.60 |
| Non-smoker vs Smoker | 16 | 46 | 3.21E-05 | 2.13 |
| Type 2 Diabetes | 5 | 16 | 5.99E-05 | 4.24 |
| Early Disseminated | 1 | 12 | 1.11E-04 | 5.35 |
| Immune response | 46 | 83 | 1.20E-04 | 1.59 |
| Cigarette smoking | 36 | 61 | 1.57E-04 | 1.77 |
| Monocytes | 10 | 34 | 2.21E-04 | 2.26 |
| Airway epithelium | 26 | 58 | 2.21E-04 | 1.78 |
| Reactive oxygen species | 12 | 58 | 2.69E-04 | 1.76 |
| Mycobacterium tuberculosis | 3 | 14 | 3.67E-04 | 3.97 |
| Smoking Status: Current, Never | 23 | 36 | 6.22E-04 | 2.08 |
| Chronic obstructive pulmonary disease | 16 | 30 | 6.36E-04 | 2.26 |
| Polymorphonuclear leukocytes | 10 | 55 | 1.08E-03 | 1.70 |
| Anaplasma phagocytophilum | 10 | 55 | 1.08E-03 | 1.70 |
| Granulocytic anaplasmosis | 10 | 55 | 1.08E-03 | 1.70 |
| Metabolism | 7 | 35 | 1.27E-03 | 2.01 |
| Epithelial gene expression | 16 | 36 | 1.27E-03 | 1.98 |
| Lyme disease | 2 | 15 | 1.27E-03 | 3.24 |
| Borrelia burgdorferi | 2 | 15 | 1.27E-03 | 3.24 |
| PBMCs | 22 | 58 | 1.27E-03 | 1.65 |
| DE genes expressed in Obese, Lean | 3 | 94 | 2.28E-03 | 1.39 |
| Obese vs Lean | 2 | 94 | 2.28E-03 | 1.39 |
| Apoptosis | 34 | 85 | 4.29E-03 | 1.40 |
| Protein catabolism | 2 | 10 | 5.26E-03 | 3.63 |
| Plasmodium falciparum | 1 | 14 | 5.26E-03 | 2.83 |
| Malaria | 1 | 14 | 5.26E-03 | 2.83 |
| Blood monocytes | 1 | 14 | 5.26E-03 | 2.83 |
| Hepatocellular carcinoma | 1 | 29 | 5.26E-03 | 1.95 |
| HBV Infection | 1 | 29 | 5.26E-03 | 1.95 |
| Infection type: Chronic | 29 | 66 | 5.60E-03 | 1.47 |
| Infection induced gene expression | 110 | 148 | 6.10E-03 | 1.21 |
| Pack-years: (10 - 20) | 5 | 14 | 6.83E-03 | 2.74 |
| Diet intake: Dietary energy restriction | 3 | 25 | 7.21E-03 | 2.01 |
| DE genes expressed in Obese | 15 | 33 | 7.27E-03 | 1.80 |
| Idiopathic pulmonary fibrosis | 1 | 13 | 7.33E-03 | 2.82 |
| Cytokines | 4 | 12 | 1.00E-02 | 2.83 |
| Lung cancer | 8 | 23 | 1.00E-02 | 2.01 |
| Viral response | 9 | 25 | 1.00E-02 | 1.94 |
| Mannose metabolism | 1 | 88 | 1.00E-02 | 1.34 |
| Insulin resistance | 7 | 89 | 1.00E-02 | 1.33 |
| Adipose tissue gene expression | 3 | 88 | 1.05E-02 | 1.33 |
| DE genes expressed in Healthy | 11 | 31 | 1.09E-02 | 1.77 |
| Before vs After diet intake | 7 | 19 | 1.23E-02 | 2.13 |
| Blood immune cells | 20 | 37 | 1.23E-02 | 1.65 |
| Influenza A Infection | 20 | 37 | 1.23E-02 | 1.65 |
| E. coli infection | 20 | 37 | 1.23E-02 | 1.65 |
| Staphylococcus aureus infection | 20 | 37 | 1.23E-02 | 1.65 |
| Streptococcus pneumoniae infection | 20 | 37 | 1.23E-02 | 1.65 |
| T effector cells | 2 | 11 | 1.39E-02 | 2.82 |
| Helminth infection | 2 | 11 | 1.39E-02 | 2.82 |
| Macrophages | 17 | 49 | 1.53E-02 | 1.50 |
| Lipid metabolism | 9 | 33 | 1.58E-02 | 1.68 |
| Infection induced gene expression in mice | 18 | 39 | 1.65E-02 | 1.58 |
| Dendritic cells | 20 | 73 | 1.65E-02 | 1.35 |
| Vascularization | 1 | 13 | 1.77E-02 | 2.42 |
| Energy restriction associated gene expression | 2 | 22 | 1.77E-02 | 1.90 |
| Oxidative stress | 11 | 24 | 1.89E-02 | 1.82 |
| Hematopoiesis | 2 | 13 | 1.95E-02 | 2.38 |
| Vesicular traffic | 1 | 12 | 2.10E-02 | 2.43 |
| DE genes expressed in Insulin sensitive individuals | 1 | 12 | 2.10E-02 | 2.43 |
| Lipid induced gene expression | 1 | 15 | 2.22E-02 | 2.15 |
| Xenobiotic metabolism | 4 | 13 | 2.66E-02 | 2.26 |
| Bacterial infection | 4 | 13 | 2.82E-02 | 2.24 |
| Protein Metabolism | 2 | 12 | 3.00E-02 | 2.31 |
| Skeletal muscle gene expression | 2 | 12 | 3.03E-02 | 2.30 |
| Maternal cigarette smoking | 2 | 13 | 3.03E-02 | 2.20 |
| Mosquito-borne pathogen | 7 | 21 | 3.58E-02 | 1.77 |
| Signal Transduction | 7 | 15 | 3.62E-02 | 2.01 |
| Zika virus | 8 | 21 | 4.12E-02 | 1.75 |
| Pack-years: (20 - 30; Healthy smoker), (Above 40; Smoker with COPD) | 4 | 12 | 4.50E-02 | 2.15 |

**Table 3.** Collation of results obtained on query of E.PAGE with genes differentially expressed in Rheumatoid Arthritis.

| Gene Modules | Number of Modules | Number of DE Genes | P<sub>adj</sub> | Fold Enrichment |
| --- | --- | --- | --- | --- |
| Infection type: Acute | 58 | 131 | 8.49E-34 | 2.63 |
| Immune response | 46 | 107 | 1.03E-27 | 2.87 |
| Infection induced gene expression | 110 | 159 | 1.03E-27 | 1.82 |
| Inflammation | 31 | 91 | 5.26E-22 | 2.88 |
| PBMCs | 22 | 78 | 6.89E-20 | 3.10 |
| Transcription regulation | 11 | 44 | 1.16E-19 | 5.77 |
| Interferons | 15 | 63 | 3.98E-19 | 3.70 |
| Central nervous system | 4 | 27 | 4.96E-18 | 10.49 |
| Infection type: Chronic | 29 | 85 | 5.19E-18 | 2.65 |
| Astrocytes | 2 | 17 | 1.94E-17 | 24.50 |
| Plasmodium falciparum | 1 | 29 | 9.43E-17 | 8.21 |
| Malaria | 1 | 29 | 9.43E-17 | 8.21 |
| Blood monocytes | 1 | 29 | 9.43E-17 | 8.21 |
| Dendritic cells | 20 | 91 | 1.85E-16 | 2.36 |
| Mycobacterium tuberculosis | 3 | 25 | 2.53E-16 | 9.93 |
| Infection induced gene expression in mice | 18 | 59 | 4.97E-16 | 3.35 |
| Pro-inflammatory response | 1 | 15 | 1.72E-15 | 24.29 |
| Chemokines | 2 | 17 | 3.11E-15 | 17.76 |
| Viral response | 9 | 41 | 9.93E-15 | 4.46 |
| Monocytes | 10 | 43 | 7.69E-14 | 4.00 |
| Olive oil induced gene expression | 3 | 17 | 2.43E-13 | 13.61 |
| Bacterial infection | 4 | 27 | 2.79E-13 | 6.51 |
| Dyslipemia | 1 | 14 | 3.35E-13 | 19.40 |
| Bone marrow monocytes | 1 | 16 | 3.35E-13 | 14.78 |
| Myelodysplastic syndromes | 1 | 16 | 3.35E-13 | 14.78 |
| Hematopoietic stem cell disease | 1 | 16 | 3.35E-13 | 14.78 |
| Lyme disease | 2 | 24 | 7.14E-13 | 7.25 |
| Borrelia burgdorferi | 2 | 24 | 7.14E-13 | 7.25 |
| IL-12 | 4 | 39 | 4.66E-12 | 3.84 |
| Th1-mediated response | 4 | 39 | 4.66E-12 | 3.84 |
| Parasite killing | 4 | 39 | 4.66E-12 | 3.84 |
| Diet intake: Olive oil | 2 | 15 | 7.67E-12 | 13.43 |
| Airway epithelium | 26 | 60 | 2.08E-11 | 2.57 |
| DE genes expressed in Obese | 15 | 43 | 3.55E-11 | 3.28 |
| Blood immune cells | 20 | 48 | 3.55E-11 | 2.99 |
| Influenza A Infection | 20 | 48 | 3.55E-11 | 2.99 |
| E. coli infection | 20 | 48 | 3.55E-11 | 2.99 |
| Staphylococcus aureus infection | 20 | 48 | 3.55E-11 | 2.99 |
| Streptococcus pneumoniae infection | 20 | 48 | 3.55E-11 | 2.99 |
| Mosquito-borne pathogen | 7 | 34 | 4.44E-11 | 4.02 |
| Zika virus | 8 | 34 | 6.48E-11 | 3.96 |
| Tissue remodeling | 1 | 10 | 1.04E-09 | 19.74 |
| Immunoregulation | 1 | 10 | 1.04E-09 | 19.74 |
| Hepatocellular carcinoma | 1 | 36 | 1.13E-09 | 3.38 |
| HBV Infection | 1 | 36 | 1.13E-09 | 3.38 |
| Chronic obstructive pulmonary disease | 16 | 33 | 3.61E-09 | 3.48 |
| Lipid metabolism | 9 | 41 | 3.98E-09 | 2.91 |
| Cigarette smoking | 36 | 57 | 4.61E-09 | 2.31 |
| Macrophages | 17 | 54 | 1.51E-08 | 2.31 |
| HIV infection | 9 | 30 | 1.59E-08 | 3.54 |
| Non-smoker vs Smoker | 16 | 42 | 1.60E-08 | 2.72 |
| Metabolism | 7 | 37 | 1.82E-08 | 2.97 |
| Zika virus associated pDCs response | 1 | 15 | 3.13E-08 | 7.21 |
| Early Disseminated | 1 | 13 | 8.80E-08 | 8.11 |
| Apoptosis | 34 | 78 | 9.83E-08 | 1.80 |
| Type 2 Diabetes | 5 | 16 | 1.40E-07 | 5.93 |
| Reactive oxygen species | 12 | 52 | 1.44E-07 | 2.21 |
| Fusobacterium nucleatum | 3 | 10 | 1.58E-07 | 11.53 |
| Oral pathogen | 3 | 10 | 1.58E-07 | 11.53 |
| Diet intake: Low calorie diet | 4 | 23 | 1.60E-07 | 3.98 |
| Epithelial gene expression | 16 | 36 | 1.63E-07 | 2.77 |
| Smoking Status: Current, Never | 23 | 35 | 1.67E-07 | 2.82 |
| DE genes expressed in Healthy | 11 | 35 | 1.97E-07 | 2.80 |
| Echovirus-30 | 1 | 13 | 2.33E-07 | 7.35 |
| Blood,ÄCerebrospinal Fluid Barrier | 1 | 13 | 2.33E-07 | 7.35 |
| Polar Infection | 1 | 13 | 2.33E-07 | 7.35 |
| Skeletal muscle gene expression | 2 | 18 | 3.23E-07 | 4.83 |
| Before vs After diet intake | 7 | 23 | 8.32E-07 | 3.60 |
| Pack-years: (20 - 30; Healthy smoker), (Above 40; Smoker with COPD) | 4 | 18 | 8.72E-07 | 4.50 |
| T effector cells | 2 | 15 | 1.08E-06 | 5.38 |
| Helminth infection | 2 | 15 | 1.08E-06 | 5.38 |
| Cell growth | 7 | 15 | 1.37E-06 | 5.27 |
| Macrophages gene expression | 4 | 12 | 1.40E-06 | 6.89 |
| Cell culture based smoking effect | 3 | 13 | 1.62E-06 | 6.12 |
| Cystic Fibrosis | 1 | 24 | 1.62E-06 | 3.33 |
| Rhinovirus infection | 1 | 24 | 1.62E-06 | 3.33 |
| Human choroid plexus epithelial cells | 1 | 12 | 1.79E-06 | 6.70 |
| Cytokines | 4 | 15 | 2.70E-06 | 4.96 |
| SIV infection | 6 | 24 | 2.81E-06 | 3.22 |
| Weight associated gene expression | 10 | 16 | 2.90E-06 | 4.59 |
| Polymorphonuclear leukocytes | 10 | 48 | 2.90E-06 | 2.07 |
| Anaplasma phagocytophilum | 10 | 48 | 2.90E-06 | 2.07 |
| Granulocytic anaplasmosis | 10 | 48 | 2.90E-06 | 2.07 |
| Ulcerative colitis | 1 | 10 | 3.48E-06 | 7.92 |
| Crohn's disease | 1 | 10 | 3.48E-06 | 7.92 |
| Jurkat cells gene expression | 1 | 10 | 3.48E-06 | 7.92 |
| Pack-years: (10 - 20) | 5 | 16 | 5.00E-06 | 4.38 |
| Diet intake: Dietary energy restriction | 3 | 26 | 5.00E-06 | 2.92 |
| Viral infection | 19 | 62 | 6.62E-06 | 1.78 |
| Signal Transduction | 7 | 19 | 9.18E-06 | 3.57 |
| Vesicular traffic | 1 | 15 | 1.32E-05 | 4.26 |
| DE genes expressed in Insulin sensitive individuals | 1 | 15 | 1.32E-05 | 4.26 |
| Protein Metabolism | 2 | 15 | 2.30E-05 | 4.04 |
| Idiopathic pulmonary fibrosis | 1 | 14 | 2.58E-05 | 4.26 |
| Lung cancer | 8 | 23 | 3.20E-05 | 2.81 |
| Oxidative stress | 11 | 25 | 3.26E-05 | 2.66 |
| Non-smoker vs Smoker (Healthy smoker, Smoker with COPD) | 11 | 18 | 4.67E-05 | 3.26 |
| Zika virus associated CD4T cell response | 1 | 10 | 1.23E-04 | 5.06 |
| Diet intake vs Control | 17 | 19 | 1.64E-04 | 2.83 |
| Cytoskeletal function | 3 | 21 | 1.72E-04 | 2.64 |
| Pathogen sensing | 6 | 14 | 3.90E-04 | 3.23 |
| Antimicrobial defense | 6 | 14 | 3.90E-04 | 3.23 |
| Suppression of T cell activation | 6 | 14 | 3.90E-04 | 3.23 |
| Enhanced bactericidal activity | 6 | 14 | 3.90E-04 | 3.23 |
| Inhibition of granuloma destruction | 6 | 14 | 3.90E-04 | 3.23 |
| Viral responses | 4 | 11 | 4.20E-04 | 3.90 |
| Genotoxic | 2 | 14 | 4.28E-04 | 3.19 |
| Carcinogen | 2 | 14 | 4.28E-04 | 3.19 |
| Chemical induced gene expression | 3 | 14 | 5.13E-04 | 3.13 |
| Energy restriction associated gene expression | 2 | 20 | 7.40E-04 | 2.41 |
| Calorie restriction effect on old vs young | 1 | 14 | 1.07E-03 | 2.90 |
| Innate Immunity | 5 | 46 | 1.11E-03 | 1.63 |
| Regulatory T cells | 2 | 10 | 1.38E-03 | 3.60 |
| Immunopathology | 2 | 10 | 1.38E-03 | 3.60 |
| Helminth Infection | 2 | 10 | 1.38E-03 | 3.60 |
| Insulin resistance | 7 | 67 | 2.63E-03 | 1.40 |
| Diet intake: High-fat | 13 | 19 | 3.42E-03 | 2.16 |
| Non-genotoxic | 1 | 12 | 3.47E-03 | 2.79 |
| Hepatocarcinogens | 1 | 12 | 3.47E-03 | 2.79 |
| Liver-based in vitro models | 1 | 12 | 3.47E-03 | 2.79 |
| Immune response | 2 | 36 | 4.56E-03 | 1.64 |
| Dendritic cell maturation | 2 | 36 | 4.56E-03 | 1.64 |
| Newcastle disease virus | 2 | 36 | 4.56E-03 | 1.64 |
| Adipose tissue gene expression | 3 | 64 | 7.85E-03 | 1.36 |
| Mannose metabolism | 1 | 63 | 1.14E-02 | 1.34 |
| Hematogenous dissemination of virus | 6 | 15 | 1.54E-02 | 2.04 |
| Epidermal growth factor receptor/PI3K signaling pathway | 6 | 15 | 1.54E-02 | 2.04 |
| Obese vs Lean | 2 | 63 | 1.89E-02 | 1.31 |
| DE genes expressed in Obese, Lean | 3 | 63 | 1.90E-02 | 1.31 |
| Lipid induced gene expression | 1 | 11 | 2.42E-02 | 2.21 |
| CD4+ T cell | 7 | 11 | 2.53E-02 | 2.19 |
| Pack-years: (20 - 30) | 9 | 16 | 3.38E-02 | 1.80 |

### Case study 3: Regulation of the cell-cycle process in small cell lung cancer

We next studied gene modules associated with small cell lung cancer. The query signature containing 71 DE genes was derived from 23 clinical small cell lung cancer samples and 42 healthy control tissues (13). We found that several lungs cancer associated gene modules were infections were was the most common environmental factor associated with the DE genes statistically significant (Table 4). The effect of Cigarette smoking (*Tumor tissue vs Non tumor tissue in Non-smoker vs Smoker, Cigarette smoking, Smoking Status: Current, Never*) was also evident. As expected, *Lung tissue gene expression* and *Adenocarcinoma* were amongst the top five gene modules, along with *Cytoprotective mechanism, Mitotic spindle formation genes* and *Cell cycle*, which are important pathways dysregulated in cancer (Table 4). Other interesting gene modules that are known to be involved in lung cancer were also identified, including *Lung cancer, Cigarette smoking, Airway epithelium* and *Immune response*.

**Table 4.** Collation of results obtained on query of E.PAGE with genes differentially expressed in small cell lung cancer.

| Gene Modules | Number of Modules | Number of DE Genes | P<sub>adj</sub> | Fold Enrichment |
| --- | --- | --- | --- | --- |
| Cytoprotective mechanism | 1 | 21 | 7.28E-08 | 5.33 |
| Mitotic spindle formation genes | 1 | 10 | 1.95E-07 | 15.80 |
| Cell cycle | 4 | 10 | 8.49E-07 | 12.98 |
| Lungs tissue gene expression | 2 | 10 | 8.49E-07 | 12.62 |
| Adenocarcinoma | 2 | 10 | 3.62E-04 | 6.39 |
| Tumor tissue vs Non tumor tissue in Non-smoker vs Smoker | 3 | 10 | 6.78E-04 | 5.82 |
| Apoptosis | 34 | 34 | 9.36E-04 | 1.97 |
| Smoking Status: Current, Former, Never | 5 | 10 | 1.21E-03 | 5.27 |
| Reactive oxygen species | 12 | 23 | 1.31E-03 | 2.47 |
| Polymorphonuclear leukocytes | 10 | 22 | 2.30E-03 | 2.39 |
| Anaplasma phagocytophilum | 10 | 22 | 2.30E-03 | 2.39 |
| Granulocytic anaplasmosis | 10 | 22 | 2.30E-03 | 2.39 |
| Cigarette smoking | 36 | 23 | 2.30E-03 | 2.35 |
| Macrophages | 17 | 22 | 2.38E-03 | 2.38 |
| Smoking Status: Current, Never | 23 | 14 | 1.10E-02 | 2.85 |
| Infection induced gene expression in mice | 18 | 17 | 1.35E-02 | 2.44 |
| Lung cancer | 8 | 10 | 4.05E-02 | 3.08 |

### Case study 4: Genotoxicity associated with cobalt exposed gene expression

We next used E.PAGE to understand the gene expression pathways involved in cobalt exposure. We used 27 DE genes uncovered by measuring the effect of cobalt exposure on gene expression in two rat liver derived cell lines using microarray analysis (14). Cobalt exposed DE genes were associated with chemical induced gene expression. Other significant gene modules include *genotoxicity, carcinogen, non-genotoxic, hepatocarcinogens*, and *liver-based in vitro models* (Table 5).

**Table 5.** Collation of results obtained on query of E.PAGE with genes differentially expressed in cobalt exposure.

| Gene Modules | Number of Modules | Number of DE Genes | P<sub>adj</sub> | Fold Enrichment |
| --- | --- | --- | --- | --- |
| Genotoxic | 2 | 8 | 1.84E-05 | 12.09 |
| Non-genotoxic | 1 | 8 | 1.84E-05 | 12.33 |
| Carcinogen | 2 | 8 | 1.84E-05 | 12.09 |
| Hepatocarcinogens | 1 | 8 | 1.84E-05 | 12.33 |
| Liver-based in vitro models | 1 | 8 | 1.84E-05 | 12.33 |
| Chemical induced gene expression | 3 | 8 | 1.84E-05 | 11.87 |

### Case study 5: Single-cell COVID-19 dataset

From a single-cell RNA sequencing dataset (15), we first conducted a standard Seurat pipeline to determine the graph based clusters (18). We then analysed enrichment of gene modules based on DE genes in Seurat clusters in COVID-19 and healthy cases. As expected, we identified *COVID-19, SARS-COV2* modules. Significant enrichment was also observed for the *Inflammation, Infection-type: Acute, Immune response, Infection induced gene expression* and *Cigarette smoking* amongst the top modules that were previously shown to be COVID-19-related (15, 19, 20) (Table 6).

**Table 6.** Collation of results obtained on querying E.PAGE with genes differentially expressed in severe COVID-19.

| Gene Modules | Number of Modules | Number of DE Genes | P<sub>adj</sub> | Fold Enrichment |
| --- | --- | --- | --- | --- |
| Inflammation | 31 | 188 | 1.18E-60 | 3.39 |
| Infection type: Acute | 58 | 225 | 1.02E-56 | 2.58 |
| Immune response | 46 | 187 | 7.61E-49 | 2.86 |
| Infection induced gene expression | 110 | 273 | 1.41E-44 | 1.79 |
| Interferons | 15 | 123 | 1.89E-43 | 4.11 |
| Cigarette smoking | 36 | 143 | 6.85E-41 | 3.31 |
| Chronic obstructive pulmonary disease | 16 | 90 | 1.80E-39 | 5.42 |
| PBMCs | 22 | 139 | 3.93E-37 | 3.15 |
| DE genes expressed in Obese | 15 | 99 | 2.19E-35 | 4.30 |
| Mycobacterium tuberculosis | 3 | 49 | 9.60E-35 | 11.09 |
| Non-smoker vs Smoker | 16 | 105 | 4.88E-34 | 3.88 |
| Infection type: Chronic | 29 | 151 | 3.17E-33 | 2.69 |
| Monocytes | 10 | 87 | 6.19E-33 | 4.61 |
| Macrophages | 17 | 126 | 5.42E-32 | 3.08 |
| IL-12 | 4 | 83 | 1.20E-31 | 4.66 |
| Th1-mediated response | 4 | 83 | 1.20E-31 | 4.66 |
| Parasite killing | 4 | 83 | 1.20E-31 | 4.66 |
| Viral response | 9 | 78 | 1.05E-30 | 4.84 |
| Macrophages gene expression | 4 | 39 | 4.60E-30 | 12.77 |
| Lung cancer | 8 | 73 | 5.74E-30 | 5.09 |
| Mosquito-borne pathogen | 7 | 74 | 6.99E-30 | 4.99 |
| Zika virus | 8 | 74 | 1.72E-29 | 4.92 |
| Reactive oxygen species | 12 | 121 | 1.30E-28 | 2.93 |
| Diet intake: Dietary energy restriction | 3 | 74 | 1.55E-28 | 4.74 |
| Airway epithelium | 26 | 118 | 4.15E-27 | 2.88 |
| Plasmodium falciparum | 1 | 46 | 3.05E-25 | 7.43 |
| Malaria | 1 | 46 | 3.05E-25 | 7.43 |
| Blood monocytes | 1 | 46 | 3.05E-25 | 7.43 |
| Metabolism | 7 | 82 | 4.91E-25 | 3.76 |
| Pack-years: (10 - 20) | 5 | 46 | 1.16E-24 | 7.19 |
| Polymorphonuclear leukocytes | 10 | 112 | 7.05E-24 | 2.76 |
| Anaplasma phagocytophilum | 10 | 112 | 7.05E-24 | 2.76 |
| Granulocytic anaplasmosis | 10 | 112 | 7.05E-24 | 2.76 |
| Bone marrow monocytes | 1 | 28 | 1.65E-23 | 14.75 |
| Myelodysplastic syndromes | 1 | 28 | 1.65E-23 | 14.75 |
| Hematopoietic stem cell disease | 1 | 28 | 1.65E-23 | 14.75 |
| Apoptosis | 34 | 159 | 4.18E-23 | 2.09 |
| Energy restriction associated gene expression | 2 | 64 | 8.15E-23 | 4.40 |
| Idiopathic pulmonary fibrosis | 1 | 42 | 8.63E-23 | 7.28 |
| Smoking Status: Current, Never | 23 | 78 | 1.41E-22 | 3.59 |
| Epithelial gene expression | 16 | 79 | 5.50E-22 | 3.47 |
| Dendritic cells | 20 | 144 | 5.52E-21 | 2.13 |
| Lyme disease | 2 | 40 | 7.47E-21 | 6.89 |
| Borrelia burgdorferi | 2 | 40 | 7.47E-21 | 6.89 |
| Hepatocellular carcinoma | 1 | 69 | 1.70E-20 | 3.69 |
| HBV Infection | 1 | 69 | 1.70E-20 | 3.69 |
| Blood immune cells | 20 | 85 | 5.45E-20 | 3.02 |
| Influenza A Infection | 20 | 85 | 5.45E-20 | 3.02 |
| E. coli infection | 20 | 85 | 5.45E-20 | 3.02 |
| Staphylococcus aureus infection | 20 | 85 | 5.45E-20 | 3.02 |
| Streptococcus pneumoniae infection | 20 | 85 | 5.45E-20 | 3.02 |
| Chemokines | 2 | 24 | 5.79E-20 | 14.29 |
| Central nervous system | 4 | 35 | 5.94E-20 | 7.75 |
| Zika virus associated pDCs response | 1 | 32 | 7.36E-20 | 8.77 |
| Pack-years: (20 - 30; Healthy smoker), (Above 40; Smoker with COPD) | 4 | 42 | 1.05E-19 | 5.99 |
| Tissue remodeling | 1 | 19 | 1.94E-19 | 21.39 |
| Immunoregulation | 1 | 19 | 1.94E-19 | 21.39 |
| Sepsis | 1 | 17 | 7.12E-19 | 25.40 |
| CD14+ Monocytes | 1 | 17 | 7.12E-19 | 25.40 |
| Innate immune response | 1 | 17 | 7.12E-19 | 25.40 |
| Fatty acid metabolism | 3 | 17 | 1.15E-16 | 19.41 |
| Non-smoker vs Smoker (Healthy smoker, Smoker with COPD) | 11 | 44 | 4.18E-16 | 4.54 |
| Bacterial infection | 4 | 38 | 7.17E-16 | 5.22 |
| Early Disseminated | 1 | 25 | 9.19E-16 | 8.89 |
| Bronchoalveolar epithelium | 1 | 13 | 1.04E-14 | 26.06 |
| Olive oil induced gene expression | 3 | 21 | 5.82E-14 | 9.59 |
| HIV infection | 9 | 51 | 8.54E-14 | 3.44 |
| SARS-COV2 | 3 | 18 | 5.31E-13 | 10.88 |
| COVID-19 | 3 | 18 | 5.31E-13 | 10.88 |
| Infection induced gene expression in mice | 18 | 76 | 5.83E-13 | 2.46 |
| Astrocytes | 2 | 16 | 6.21E-13 | 13.15 |
| Citric acid cycle | 1 | 13 | 8.34E-13 | 19.08 |
| Complement system | 1 | 13 | 8.34E-13 | 19.08 |
| Diet intake: Milk fat and protein | 1 | 13 | 8.34E-13 | 19.08 |
| Apoptosis | 1 | 13 | 3.39E-12 | 17.23 |
| Human gingival fibroblasts | 2 | 13 | 4.17E-12 | 16.96 |
| Transcription regulation | 11 | 45 | 6.79E-12 | 3.37 |
| Diet intake: Olive oil | 2 | 18 | 9.01E-12 | 9.19 |
| Oxidative stress | 11 | 50 | 1.57E-11 | 3.03 |
| Dyslipemia | 1 | 15 | 1.60E-11 | 11.85 |
| Fusobacterium nucleatum | 3 | 16 | 1.89E-11 | 10.52 |
| Oral pathogen | 3 | 16 | 1.89E-11 | 10.52 |
| Pro-inflammatory response | 1 | 14 | 2.47E-11 | 12.93 |
| Atherosclerosis | 1 | 10 | 2.67E-11 | 24.91 |
| Atherosclerotic cardiovascular disease (ASCVD) | 1 | 10 | 2.67E-11 | 24.91 |
| Aging | 1 | 10 | 2.67E-11 | 24.91 |
| T effector cells | 2 | 26 | 3.15E-11 | 5.32 |
| Helminth infection | 2 | 26 | 3.15E-11 | 5.32 |
| Smoking Status: Current, Former, Never | 5 | 34 | 3.20E-11 | 4.05 |
| Oxidative phosphorylation | 3 | 13 | 2.25E-10 | 12.42 |
| Tumor tissue vs Non tumor tissue in Non-smoker vs Smoker | 3 | 31 | 2.34E-10 | 4.08 |
| Xenobiotic metabolism | 4 | 30 | 3.13E-10 | 4.16 |
| Human choroid plexus epithelial cells | 1 | 20 | 3.73E-10 | 6.37 |
| Adenocarcinoma | 2 | 29 | 5.47E-10 | 4.19 |
| Pack-years: (20 - 30) | 9 | 45 | 8.40E-10 | 2.89 |
| Regulatory T cells | 2 | 24 | 8.95E-10 | 4.93 |
| Immunopathology | 2 | 24 | 8.95E-10 | 4.93 |
| Helminth Infection | 2 | 24 | 8.95E-10 | 4.93 |
| Cell culture based smoking effect | 3 | 21 | 1.12E-09 | 5.64 |
| Hematopoiesis | 2 | 28 | 1.92E-09 | 4.09 |
| Signal Transduction | 7 | 33 | 2.01E-09 | 3.54 |
| Cystic Fibrosis | 1 | 39 | 2.33E-09 | 3.09 |
| Rhinovirus infection | 1 | 39 | 2.33E-09 | 3.09 |
| Angiogenesis | 2 | 14 | 2.35E-09 | 9.13 |
| Extracellular matrix metabolism | 1 | 10 | 3.66E-09 | 15.51 |
| Autosomal-dominant hyper-IgE syndrome | 1 | 10 | 3.66E-09 | 15.51 |
| Immunodeficiency | 1 | 10 | 3.66E-09 | 15.51 |
| Lipid metabolism | 9 | 58 | 3.95E-09 | 2.35 |
| Vascularization | 1 | 27 | 5.71E-09 | 4.01 |
| Oxidant-related | 2 | 13 | 9.38E-09 | 9.13 |
| Zika virus associated mDCs response | 1 | 19 | 2.74E-08 | 5.21 |
| Maternal cigarette smoking | 2 | 27 | 3.99E-08 | 3.65 |
| Cell death | 1 | 20 | 8.38E-08 | 4.60 |
| Leptin resistance | 1 | 11 | 9.12E-08 | 9.62 |
| Weight loss | 2 | 11 | 3.15E-07 | 8.53 |
| Gene expression induced due to fasting | 3 | 13 | 3.26E-07 | 6.76 |
| Diet intake: Fasting | 3 | 13 | 3.26E-07 | 6.76 |
| DE genes expressed in Healthy | 11 | 49 | 3.99E-07 | 2.24 |
| Cytokines | 4 | 21 | 4.55E-07 | 3.96 |
| Diet intake: Low calorie diet | 4 | 30 | 5.45E-07 | 2.96 |
| SIV infection | 6 | 35 | 5.45E-07 | 2.68 |
| Zika virus associated CD8T cell response | 1 | 16 | 1.24E-06 | 4.78 |
| Type 2 Diabetes | 5 | 19 | 1.44E-06 | 4.01 |
| Ulcerative colitis | 1 | 13 | 1.57E-06 | 5.87 |
| Crohn's disease | 1 | 13 | 1.57E-06 | 5.87 |
| Jurkat cells gene expression | 1 | 13 | 1.57E-06 | 5.87 |
| DNA damage | 3 | 10 | 3.48E-06 | 7.54 |
| Weight associated gene expression | 10 | 21 | 4.12E-06 | 3.44 |
| Obese vs Lean | 2 | 123 | 4.99E-06 | 1.46 |
| DE genes expressed in Obese, Lean | 3 | 123 | 5.04E-06 | 1.46 |
| Adipose tissue gene expression | 3 | 121 | 5.21E-06 | 1.46 |
| Chemical induced gene expression | 3 | 24 | 5.25E-06 | 3.06 |
| Insulin resistance | 7 | 122 | 5.28E-06 | 1.46 |
| Genotoxic | 2 | 23 | 1.27E-05 | 2.99 |
| Carcinogen | 2 | 23 | 1.27E-05 | 2.99 |
| Mannose metabolism | 1 | 119 | 1.31E-05 | 1.44 |
| Smoking History: > 19 years | 2 | 12 | 1.59E-05 | 5.14 |
| Pack-days: (1 - 1.21) | 2 | 12 | 1.59E-05 | 5.14 |
| Calorie restriction effect on old vs young | 1 | 24 | 1.87E-05 | 2.83 |
| Diet intake vs Control | 17 | 29 | 2.83E-05 | 2.46 |
| Non-genotoxic | 1 | 22 | 2.84E-05 | 2.92 |
| Hepatocarcinogens | 1 | 22 | 2.84E-05 | 2.92 |
| Liver-based in vitro models | 1 | 22 | 2.84E-05 | 2.92 |
| Cell cycle | 4 | 14 | 3.19E-05 | 4.11 |
| Zika virus associated CD4T cell response | 1 | 14 | 3.81E-05 | 4.04 |
| Viral responses | 4 | 17 | 3.81E-05 | 3.43 |
| Cigarette smoking in women | 3 | 13 | 4.07E-05 | 4.29 |
| Lungs tissue gene expression | 2 | 14 | 4.20E-05 | 4.00 |
| HIV-1 infection | 9 | 30 | 4.20E-05 | 2.36 |
| Smoking Status: Current, Former | 2 | 14 | 5.33E-05 | 3.90 |
| Tumor tissue vs Non tumor tissue in Current smoker vs Former Smoker | 2 | 14 | 5.33E-05 | 3.90 |
| Zika virus induced B cell response | 1 | 14 | 6.10E-05 | 3.84 |
| Zika virus associated B cell response | 1 | 14 | 6.10E-05 | 3.84 |
| Zika virus associated monocytes response | 1 | 14 | 6.10E-05 | 3.84 |
| Mitotic spindle formation genes | 1 | 12 | 7.90E-05 | 4.29 |
| Skeletal muscle gene expression | 2 | 19 | 1.04E-04 | 2.91 |
| Metabolic pathways | 2 | 10 | 1.64E-04 | 4.70 |
| Innate Immunity | 5 | 75 | 2.77E-04 | 1.52 |
| Pulmonary nontuberculous mycobacterial disease | 1 | 10 | 6.57E-04 | 3.93 |
| T cell signaling | 1 | 10 | 6.57E-04 | 3.93 |
| Before vs After diet intake | 7 | 24 | 1.03E-03 | 2.14 |
| Protein Metabolism | 2 | 16 | 2.17E-03 | 2.46 |
| Vesicular traffic | 1 | 15 | 3.42E-03 | 2.43 |
| DE genes expressed in Insulin sensitive individuals | 1 | 15 | 3.42E-03 | 2.43 |
| DNA Methylation | 5 | 11 | 4.73E-03 | 2.80 |
| CD4+ T cell | 7 | 17 | 1.54E-02 | 1.93 |
| Hematogenous dissemination of virus | 6 | 22 | 2.11E-02 | 1.71 |
| Epidermal growth factor receptor/PI3K signaling pathway | 6 | 22 | 2.11E-02 | 1.71 |
| Cytoskeletal function | 3 | 23 | 2.54E-02 | 1.65 |
| Cytoprotective mechanism | 1 | 27 | 3.10E-02 | 1.55 |
| Cell-adhesion | 3 | 16 | 3.67E-02 | 1.78 |
| Diet intake: High-fat | 13 | 24 | 4.03E-02 | 1.56 |

### Case study 6: Single-cell Smoking dataset

As a final case study, we attempted to identify enriched gene modules related to smoking using a single cell RNA sequencing dataset which contained data of smokers vs non-smokers (16). After processing the data using the Seurat pipeline and analyzing the single-cell expression data, gene set enrichment identified *Epithelial gene expression, Cigarette smoking, Airway epithelium*, and *Chronic obstructive pulmonary disease* as the top gene modules with highly significant p-values, confirming that smoking-related pathways were correctly predicted using E.PAGE (Table 7). Furthermore, smoking associated with gene signatures of lung-associated diseases such as *Lung cancer, Cystic fibrosis*, as well as with *Carcinogen* and respiratory infections such as *Influenza* and *COVID-19*.

**Table 7.** Collation of results obtained on querying E.PAGE with genes differentially expressed in heavy smoking subjects.

| Gene Modules | Number of Modules | Number of DE Genes | Padj | Fold Enrichment |
| --- | --- | --- | --- | --- |
| Epithelial gene expression | 16 | 198 | 3.07E-87 | 5.16 |
| Cigarette smoking | 36 | 261 | 6.38E-85 | 3.58 |
| Airway epithelium | 26 | 254 | 7.98E-85 | 3.68 |
| Non-smoker vs Smoker | 16 | 206 | 3.08E-81 | 4.52 |
| Idiopathic pulmonary fibrosis | 1 | 103 | 2.05E-73 | 10.60 |
| Chronic obstructive pulmonary disease | 16 | 158 | 2.05E-73 | 5.64 |
| Pack-years: (10 - 20) | 5 | 105 | 4.99E-71 | 9.74 |
| Lung cancer | 8 | 136 | 4.78E-62 | 5.63 |
| Smoking Status: Current, Never | 23 | 141 | 3.87E-44 | 3.85 |
| Pack-years: (20 - 30) | 9 | 113 | 3.95E-39 | 4.31 |
| Infection type: Acute | 58 | 276 | 3.54E-31 | 1.88 |
| Infection induced gene expression | 110 | 390 | 1.15E-30 | 1.51 |
| Inflammation | 31 | 207 | 2.15E-30 | 2.21 |
| Immune response | 46 | 214 | 1.04E-23 | 1.94 |
| Infection type: Chronic | 29 | 192 | 8.61E-23 | 2.03 |
| Apoptosis | 34 | 233 | 2.18E-22 | 1.82 |
| Transcription regulation | 11 | 77 | 8.42E-20 | 3.42 |
| Cystic Fibrosis | 1 | 74 | 1.82E-19 | 3.48 |
| Rhinovirus infection | 1 | 74 | 1.82E-19 | 3.48 |
| Lyme disease | 2 | 48 | 2.39E-18 | 4.91 |
| Borrelia burgdorferi | 2 | 48 | 2.39E-18 | 4.91 |
| Non-smoker vs Smoker (Healthy smoker, Smoker with COPD) | 11 | 62 | 4.36E-18 | 3.79 |
| Lipid metabolism | 9 | 106 | 4.56E-18 | 2.55 |
| Reactive oxygen species | 12 | 146 | 8.89E-18 | 2.10 |
| PBMCs | 22 | 152 | 1.48E-17 | 2.04 |
| Mycobacterium tuberculosis | 3 | 41 | 1.59E-17 | 5.51 |
| Pack-years: (20 - 30; Healthy smoker), (Above 40; Smoker with COPD) | 4 | 51 | 3.22E-17 | 4.32 |
| Macrophages | 17 | 143 | 6.22E-17 | 2.07 |
| Infection induced gene expression in mice | 18 | 119 | 6.23E-17 | 2.29 |
| Interferons | 15 | 115 | 3.24E-16 | 2.28 |
| Polymorphonuclear leukocytes | 10 | 138 | 2.89E-15 | 2.02 |
| Anaplasma phagocytophilum | 10 | 138 | 2.89E-15 | 2.02 |
| Granulocytic anaplasmosis | 10 | 138 | 2.89E-15 | 2.02 |
| Central nervous system | 4 | 38 | 6.33E-15 | 5.00 |
| Oxidative stress | 11 | 77 | 7.09E-15 | 2.77 |
| HIV infection | 9 | 72 | 9.37E-15 | 2.88 |
| Signal Transduction | 7 | 55 | 1.14E-14 | 3.50 |
| Hepatocellular carcinoma | 1 | 82 | 1.93E-14 | 2.61 |
| HBV Infection | 1 | 82 | 1.93E-14 | 2.61 |
| Human choroid plexus epithelial cells | 1 | 30 | 2.52E-13 | 5.67 |
| IL-12 | 4 | 76 | 1.01E-12 | 2.53 |
| Th1-mediated response | 4 | 76 | 1.01E-12 | 2.53 |
| Parasite killing | 4 | 76 | 1.01E-12 | 2.53 |
| Monocytes | 10 | 78 | 2.14E-12 | 2.45 |
| Dendritic cells | 20 | 186 | 3.07E-12 | 1.63 |
| Smoking Status: Current, Former | 2 | 30 | 7.44E-12 | 4.96 |
| Tumor tissue vs Non tumor tissue in Current smoker vs Former Smoker | 2 | 30 | 7.44E-12 | 4.96 |
| Bronchoalveolar epithelium | 1 | 13 | 1.11E-11 | 15.46 |
| Viral response | 9 | 69 | 1.16E-11 | 2.54 |
| Squamous cell lung carcinoma | 1 | 26 | 2.11E-11 | 5.51 |
| Smoking Years Quit: > 2 years | 1 | 26 | 2.11E-11 | 5.51 |
| Pack-years: (30 - 40) | 1 | 26 | 2.11E-11 | 5.51 |
| Metabolism | 7 | 82 | 6.06E-11 | 2.23 |
| Cytoprotective mechanism | 1 | 70 | 1.35E-10 | 2.38 |
| Mosquito-borne pathogen | 7 | 63 | 1.57E-10 | 2.52 |
| Zika virus associated pDCs response | 1 | 28 | 2.84E-10 | 4.55 |
| Zika virus | 8 | 63 | 2.84E-10 | 2.48 |
| SARS-COV2 | 3 | 19 | 4.46E-10 | 6.81 |
| COVID-19 | 3 | 19 | 4.46E-10 | 6.81 |
| Lungs tissue gene expression | 2 | 27 | 5.51E-10 | 4.57 |
| DE genes expressed in Obese | 15 | 82 | 7.48E-10 | 2.11 |
| Mucus overproduction | 2 | 18 | 8.21E-10 | 7.02 |
| Skeletal muscle gene expression | 2 | 37 | 1.35E-09 | 3.36 |
| Cell culture based smoking effect | 3 | 27 | 2.03E-09 | 4.30 |
| Obese vs Lean | 2 | 208 | 2.03E-09 | 1.46 |
| DE genes expressed in Obese, Lean | 3 | 208 | 2.06E-09 | 1.46 |
| SIV infection | 6 | 55 | 3.91E-09 | 2.50 |
| Cytokines | 4 | 32 | 4.68E-09 | 3.58 |
| Insulin resistance | 7 | 205 | 4.75E-09 | 1.45 |
| Adipose tissue gene expression | 3 | 203 | 5.36E-09 | 1.46 |
| Mannose metabolism | 1 | 202 | 6.76E-09 | 1.45 |
| Smoking Status: Current, Former, Never | 5 | 41 | 1.02E-08 | 2.90 |
| Early Disseminated | 1 | 22 | 2.01E-08 | 4.64 |
| Blood immune cells | 20 | 88 | 6.63E-08 | 1.86 |
| Influenza A Infection | 20 | 88 | 6.63E-08 | 1.86 |
| E. coli infection | 20 | 88 | 6.63E-08 | 1.86 |
| Staphylococcus aureus infection | 20 | 88 | 6.63E-08 | 1.86 |
| Streptococcus pneumoniae infection | 20 | 88 | 6.63E-08 | 1.86 |
| Macrophages gene expression | 4 | 22 | 8.61E-08 | 4.27 |
| Mitotic spindle formation genes | 1 | 21 | 8.68E-08 | 4.45 |
| Genotoxic | 2 | 37 | 8.68E-08 | 2.86 |
| Carcinogen | 2 | 37 | 8.68E-08 | 2.86 |
| Cell cycle | 4 | 23 | 1.30E-07 | 4.01 |
| Chemical induced gene expression | 3 | 37 | 1.37E-07 | 2.80 |
| Chemokines | 2 | 16 | 1.61E-07 | 5.65 |
| Dyslipemia | 1 | 14 | 1.75E-07 | 6.57 |
| DE genes expressed in Lean | 3 | 10 | 4.35E-07 | 9.56 |
| Zika virus associated mDCs response | 1 | 23 | 4.37E-07 | 3.74 |
| Vesicular traffic | 1 | 31 | 4.60E-07 | 2.98 |
| DE genes expressed in Insulin sensitive individuals | 1 | 31 | 4.60E-07 | 2.98 |
| Protein Metabolism | 2 | 32 | 4.60E-07 | 2.92 |
| Non-genotoxic | 1 | 35 | 4.60E-07 | 2.75 |
| Hepatocarcinogens | 1 | 35 | 4.60E-07 | 2.75 |
| Liver-based in vitro models | 1 | 35 | 4.60E-07 | 2.75 |
| Astrocytes | 2 | 13 | 7.52E-07 | 6.34 |
| DE genes expressed in Healthy | 11 | 70 | 9.81E-07 | 1.90 |
| Olive oil induced gene expression | 3 | 17 | 1.08E-06 | 4.61 |
| Weight associated gene expression | 10 | 30 | 1.12E-06 | 2.91 |
| Transport | 3 | 15 | 1.18E-06 | 5.19 |
| Diet intake: Olive oil | 2 | 16 | 1.19E-06 | 4.85 |
| Diet intake: Low calorie diet | 4 | 41 | 1.30E-06 | 2.40 |
| Pro-inflammatory response | 1 | 12 | 1.39E-06 | 6.58 |
| Regulatory T cells | 2 | 26 | 1.39E-06 | 3.17 |
| Immunopathology | 2 | 26 | 1.39E-06 | 3.17 |
| Helminth Infection | 2 | 26 | 1.39E-06 | 3.17 |
| Tumor tissue vs Non tumor tissue in Non-smoker vs Smoker | 3 | 34 | 1.41E-06 | 2.66 |
| Fusobacterium nucleatum | 3 | 14 | 1.45E-06 | 5.46 |
| Oral pathogen | 3 | 14 | 1.45E-06 | 5.46 |
| Diffuse large B-cell lymphoma | 1 | 14 | 1.54E-06 | 5.42 |
| Germinal center B-cell | 1 | 14 | 1.54E-06 | 5.42 |
| DNA repair | 1 | 14 | 1.54E-06 | 5.42 |
| Genomic stability | 1 | 14 | 1.54E-06 | 5.42 |
| Prostaglandin metabolism | 1 | 10 | 3.94E-06 | 7.39 |
| DE genes expressed in Low calorie diet | 1 | 10 | 3.94E-06 | 7.39 |
| Epithelial barrier integrity | 1 | 11 | 3.94E-06 | 6.54 |
| Cilia beat activity | 1 | 11 | 3.94E-06 | 6.54 |
| Cytoskeletal function | 3 | 49 | 4.67E-06 | 2.09 |
| Oxidant-related | 2 | 12 | 2.20E-05 | 5.00 |
| Diet intake: Dietary energy restriction | 3 | 51 | 2.20E-05 | 1.94 |
| Echovirus-30 | 1 | 18 | 2.49E-05 | 3.44 |
| Blood,ÄCerebrospinal Fluid Barrier | 1 | 18 | 2.49E-05 | 3.44 |
| Polar Infection | 1 | 18 | 2.49E-05 | 3.44 |
| Adenocarcinoma | 2 | 29 | 3.16E-05 | 2.49 |
| Human papillomavirus | 2 | 11 | 4.65E-05 | 5.06 |
| Zika virus induced B cell response | 1 | 19 | 6.29E-05 | 3.09 |
| Zika virus associated B cell response | 1 | 19 | 6.29E-05 | 3.09 |
| Ulcerative colitis | 1 | 14 | 9.73E-05 | 3.75 |
| Crohn's disease | 1 | 14 | 9.73E-05 | 3.75 |
| Jurkat cells gene expression | 1 | 14 | 9.73E-05 | 3.75 |
| Viral infection | 19 | 141 | 1.10E-04 | 1.37 |
| T effector cells | 2 | 22 | 1.28E-04 | 2.67 |
| Helminth infection | 2 | 22 | 1.28E-04 | 2.67 |
| Before vs After diet intake | 7 | 38 | 1.45E-04 | 2.01 |
| Cell growth | 7 | 22 | 1.66E-04 | 2.62 |
| Innate Immunity | 5 | 116 | 3.50E-04 | 1.39 |
| Xenobiotic metabolism | 4 | 27 | 3.92E-04 | 2.22 |
| Bacterial infection | 4 | 27 | 4.44E-04 | 2.20 |
| DNA Methylation | 5 | 18 | 4.82E-04 | 2.72 |
| Energy restriction associated gene expression | 2 | 44 | 4.91E-04 | 1.80 |
| Pack-years: Above 40 | 2 | 10 | 1.14E-03 | 3.75 |
| Gene expression induced due to fasting | 3 | 11 | 1.36E-03 | 3.40 |
| Diet intake: Fasting | 3 | 11 | 1.36E-03 | 3.40 |
| Type 2 Diabetes | 5 | 19 | 1.47E-03 | 2.38 |
| Maternal cigarette smoking | 2 | 25 | 2.60E-03 | 2.01 |
| Immune reposne | 2 | 89 | 3.69E-03 | 1.37 |
| Dendritic cell maturation | 2 | 89 | 3.69E-03 | 1.37 |
| Newcastle disease virus | 2 | 89 | 3.69E-03 | 1.37 |
| Cell-adhesion | 3 | 28 | 4.32E-03 | 1.85 |
| Diet intake vs Control | 17 | 34 | 4.93E-03 | 1.71 |
| Viral responses | 4 | 18 | 5.73E-03 | 2.16 |
| Hematopoiesis | 2 | 22 | 8.83E-03 | 1.91 |
| Zika virus associated CD8T cell response | 1 | 13 | 1.22E-02 | 2.31 |
| Calorie restriction effect on old vs young | 1 | 25 | 1.33E-02 | 1.75 |
| Vascularization | 1 | 21 | 1.41E-02 | 1.85 |
| Host susceptibility | 2 | 16 | 2.10E-02 | 1.95 |
| Macrophage activation | 2 | 16 | 2.10E-02 | 1.95 |
| Inflammatory diseases | 2 | 16 | 2.10E-02 | 1.95 |
| Plasma insulin level | 5 | 12 | 2.21E-02 | 2.19 |
| Pathogen sensing | 6 | 22 | 2.31E-02 | 1.72 |
| Antimicrobial defense | 6 | 22 | 2.31E-02 | 1.72 |
| Supression of T cell activation | 6 | 22 | 2.31E-02 | 1.72 |
| Enhanced bactericidal activity | 6 | 22 | 2.31E-02 | 1.72 |
| Inhibition of granuloma destruction | 6 | 22 | 2.31E-02 | 1.72 |
| HIV-1 infection | 9 | 32 | 3.48E-02 | 1.49 |
| Plasmodium falciparum | 1 | 18 | 3.91E-02 | 1.72 |
| Malaria | 1 | 18 | 3.91E-02 | 1.72 |
| Blood monocytes | 1 | 18 | 3.91E-02 | 1.72 |
| Hematogenous dissemination of virus | 6 | 32 | 4.04E-02 | 1.47 |
| Epidermal growth factor receptor/PI3K signaling pathway | 6 | 32 | 4.04E-02 | 1.47 |

## Discussion

Environmental factors are known to influence the development of disease, with or without combination with genetic factors, however there is currently no curated database and enrichment tool to identify the genes and the corresponding biological processes associated with these environmental conditions. We developed E.PAGE, a database and enrichment tool to understand the gene-environment relationship. Our database was developed based on experimental evidence obtained from the published literature to establish a relationship between environmental factors, differentially expressed genes and specific biological processes associated with the genes.

To set up the database, we used *cigarette smoking, infections, toxic chemicals* and *diet*, as they constitute the primary environmental factors influencing disease outcomes (4). We made every effort to ensure completeness, accuracy and currency of the database. The current database has 243 datasets which consists of 25789 genes in total. The largest number of datasets relate to *diet* and *infections* due to the long research history of these two environmental factors and disease. We manually curated each dataset using specific keywords and a brief description, abstract published with these datasets. We then developed an enrichment tool that uncovers modules associated with genes of interest using the methods we previously published (10). In six case studies, we tested E.PAGE with sets of DE genes available from the literature. Specifically, we tested two gene lists associated with autoimmunity - T1D and RA - along with those related to small cell lung cancer, COVID-19 and smoking subjects. To confirm the effect of toxic chemicals on differential gene expression, we also used gene expression data from a study on cobalt exposure.

On testing T1D and RA associated DE genes, we found a large number of gene modules related to immune responses, which supports previous studies on how malfunction in the adaptive immune response results in activation of self-reactive T cells. We also obtained a substantial number of environmental modules associated with viral and bacterial infections, which supports recent findings on how bacterial and viral infections are implicated in immune response signaling in autoimmune disease pathogenesis. The T1D and RA associated DE genes were found to be primarily enriched in *infection-*associated gene modules and less in gene modules associated with the environmental factors *diet, cigarette smoking* or *toxic chemicals*. This information supports the hypotheses that infection-associated immune responses are major contributors to the development of T1D and RA (21-23). A substantial number of genes involved in the central nervous system were also related to RA, consistent with other evidence (24).

When small cell lung cancer genes were tested, we found a large number of environmental modules for DE genes to be related to *lung cancer*, as expected. We also found an expected link to *cell cycle*, since cell cycle checkpoints are disrupted leading to tumour development and cancer progression. Genes relating to *cytoprotective function, mitotic spindle formation* are also generally dysregulated in cancer. Recent studies that show a high incidence of retrovirus in lung small cell cancer suggest a possible direct link between infections and small cell cancer (25).

To further assess associations between environmental factors with toxic chemicals, we tested genes differentially expressed due to cobalt exposure against the E.PAGE database. On testing, we found the modules *Genotoxicity* and *Carcinogen* to be enriched. We also obtained a substantial number of genes differentially expressed due to toxic chemicals as environmental factors, supporting the validity of the tool to identify potential involvement of toxic chemicals on DE genes involved in critical functions in a relevant datasets.

On testing gene expression data sourced from patients with COVID-19, we found that genes differentially expressed in severe cases were linked to gene modules common between bronchoalveolar and peripheral immune environments (15, 20). This finding shows how the E.PAGE database can be used to find commonalities between two sets of differentially expressed genes, even if they may not have many genes in common.

On testing the single-cell gene expression data for smoking we found gene modules for Cigarette smoking, Airway epithelium, Epithelial gene expression, and Chronic obstructive pulmonary disease. Additional pathways that are well known to be altered by cigarette smoking were identified. Therefore, E.PAGE was able to find relevant significantly enriched gene modules.

From the above case studies, we found that our database is highly reliable and has the potential to establish a link between environmental factors and important biological processes. In the case studies, we generally obtained a higher number of DE genes related to infection as an environmental factor. Though this link with infection may be valid, there is a possibility of dataset bias due to limited type of input data such as gene list, similarities between infection and tissue damage -associated immune responses. Additionally, our study is limited to four types of environmental variables, therefore to increase usage towards wider community more environmental datasets need to be integrated.

A key benefit of this research is to predict gene-environment interactions to identify novel associations between environmental factors and disease, and to inform hypothesis synthesis and target selection. Thereby, it allows scientists and epidemiologists to dissect which genes may be influenced by environmental exposures in different disease conditions. We illustrate this by using examples from type-1 diabetes, rheumatoid arthritis, small cell lung cancer and COVID-19 datasets.

The current study lends itself to future extension to additional environmental variables such as alcohol, physical activities, life-style factors, which could facilitate developing disease risk prediction models.

## Availability and requirements

**Project name:** E.PAGE (Environmental Pathways Affecting Gene Expression)

**Project home page:** github.com/AhmedMehdiLab/E.PAGE

**Operating system(s):** Platform independent.

**Programming language:** R version 4.0.3 (www.R-project.org/)

**Other requirements**: *tidyverse* and *Seurat* R packages

## Declarations

### Ethics Approval and consent to participate

Not applicable

### Consent for publication

Not applicable

